# Using competition assays to quantitatively model cooperative binding by transcription factors and other ligands

**DOI:** 10.1101/170340

**Authors:** Jacob Peacock, James B. Jaynes

## Abstract

**BACKGROUND:** The affinities of DNA binding proteins for target sites can be used to model the regulation of gene expression. These proteins can bind to DNA cooperatively, strongly impacting their affinity and specificity. However, current methods for measuring cooperativity do not provide the means to accurately predict binding behavior over a wide range of concentrations.

**METHODS:** We use standard computational and mathematical methods, and develop novel methods as described in Results.

**RESULTS:** We explore some complexities of cooperative binding, and develop an improved method for relating *in vitro* measurements to *in vivo* function, based on ternary complex formation. We derive expressions for the equilibria among the various complexes, and explore the limitations of binding experiments that model the system using a single parameter. We describe how to use single-ligand binding and ternary complex formation in tandem to determine parameters that have thermodynamic relevance. We develop an improved method for finding both single-ligand dissociation constants and concentrations simultaneously. We show how the cooperativity factor can be found when only one of the single-protein dissociation constants can be measured.

**CONCLUSIONS:** The methods that we develop constitute an optimized approach to accurately model cooperative binding.

**GENERAL SIGNIFICANCE:** The expressions and methods we develop for modeling and analyzing DNA binding and cooperativity are applicable to most cases where multiple ligands bind to distinct sites on a common substrate. The parameters determined using these methods can be fed into models of higher-order cooperativity to increase their predictive power.

**HIGHLIGHTS:**

- Hill plots remain prominent in biology, but can mask cooperativity
- Effective modeling of binding by two ligands requires the use of 3 parameters
- We develop novel ways to find these parameters for two cooperating ligands
- We show how they can be used to enhance the power of established methods
- We describe how this framework can be extended to multiple cooperating ligands

## 1. INTRODUCTION

Cooperative binding by multiple ligands to a substrate is ubiquitous in biological systems. Methods of detecting and analyzing cooperative binding have been well developed over time at a theoretical level. Cooperative binding occurs when the binding of a first ligand to a substrate increases (or decreases) the complex’s affinity for subsequent ligands. The phenomenon was first observed and modeled in the oxygen and hemoglobin system, where the binding of one oxygen to unsaturated hemoglobin increases the affinity for the next oxygen [1]. Hill proposed a one-step cooperative binding model,

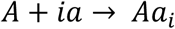

were *A* is the binding substrate, in this case hemoglobin, *a* is the ligand, oxygen, and *i* is the total number of oxygens that bind cooperatively. Using the equilibrium (association) constant for the reaction, *K_a_*, gives rise to the Hill equation for the fractional occupancy, *θ*:

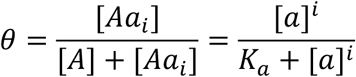

Applying the logarithm gives a linearized form, and allows determination of the Hill number, *i*, from measured [*a*] and *θ*. While computationally and experimentally accessible, the Hill model has numerous pitfalls, and a Hill plot can obscure important cooperative properties of a system (e.g., see Fig. 1 and Fig. 1C,D in Peacock and Jaynes [2]).

**Fig. 1.**
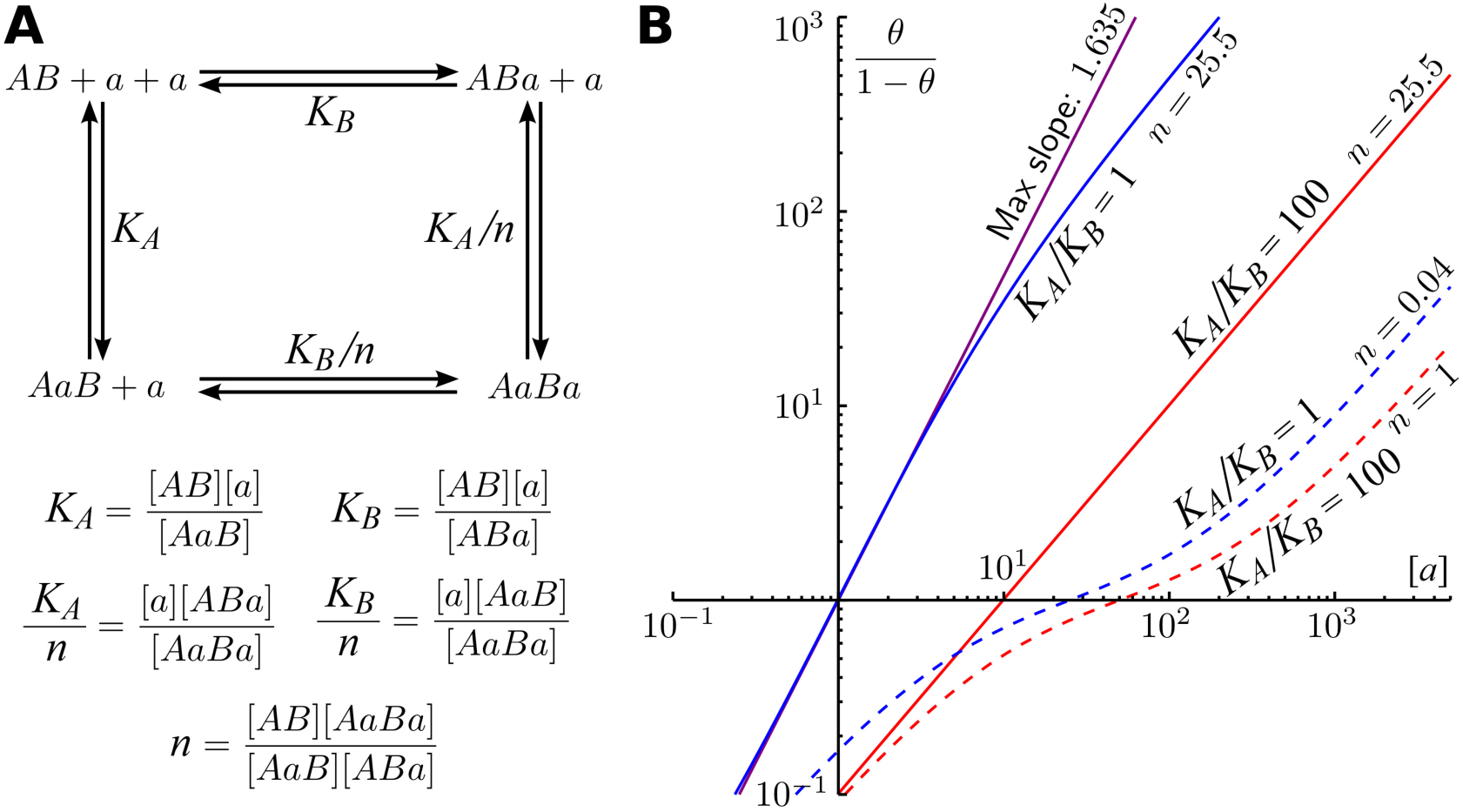
Barriers to determining cooperativity from Hill plots. <u>**A:** Model for cooperative binding</u> to *AB* (substrate with two distinct binding sites) by ligand *a*. The ternary complex *AaBa* can dissociate in two ways, losing *a* from either the *A* or the *B* site first. Defining the Kd’s for the ternary complex as *K_A_*/*n* and *K_B_*/*n* reduces the number of variables, because, from the definitions of the Kd’s (below the line), *K_A_* divided by *K_A_*/*n* gives the same thing as *K_B_* divided by *K_B_*/*n*. <u>**B.** Hill plots for two binding sites with the same or different Kd’s.</u> From the model in A, the ratio (fractional occupancy) / (1 - fractional occupancy), which is (*K_A_*+*K_B_*+2*n*[*a*])[*a*]/{2*K_A_K_B_*+(*K_A_*+*K_B_*)[*a*]} (derived in Fig. 1A of Peacock and Jaynes [2]), was used to generate Hill plots. Concentration units (for Kd’s and [*a*]) are arbitrary. The case where *K_A_* = *K_B_* = 5 and *n* = 25.5 is shown as a solid blue curve, along with a tangent line (purple) at the point of maximum slope. Also shown are: the same equivalent sites, but with negative cooperativity (*n* = 0.04, dashed blue), and the case of two non-equivalent sites (*K_A_* = 5, *K_B_* = 500), either with positive cooperativity (*n* = 25.5, solid red) or with no cooperativity (*n* = 1, dashed red). Note the similarity in shape of the plots for equivalent sites with negative cooperativity and for non-equivalent sites without cooperativity. Also note that for two non-equivalent sites, when *n* approaches the value (*K_A_*+*K_B_*)^2/4*K_A_K_B_* (derived in Fig. 1A of Peacock and Jaynes [2]), the plot approaches a straight line of slope 1, which is indistinguishable from equivalent sites with no cooperativity. Thus, without prior knowledge that sites are equivalent, Hill plots are at best ambiguous for identifying cooperativity.

The shortcomings of the Hill model in describing the hemoglobin and other systems motivated a slew of subsequent models [3], including Adair-Klotz, KNF, and MWC models [4,5]. Generally, any binding model for a given system may be expressed as a binding polynomial or partition function [3,6,7]. Any specific binding model can be related to the terms and parameters of the binding polynomial. Some authors have introduced theoretically and pedagogically useful formalism into the binding polynomial, which aid in relating the general parameters to specific binding models [8-11].

In recent decades, the importance of cooperativity in eukaryotic transcription regulation has been revealed [12-21]. Cooperativity among transcription factors and cofactors is crucial for achieving nucleic acid binding specificity *in vivo*, allowing a relatively small group of transcription factors to combinatorially regulate large genomes (see e.g. [12]). Using *in vitro* approaches such as gel shifts (EMSA, or electromobility shift analysis) of labeled oligonucleotides (oligos) by purified binding proteins and surface plasmon resonance (SPR), cooperativity has been revealed and quantified in a few cases, serving as conceptual models [22-29]. While SPR can provide more detailed interaction information than EMSAs, it is limited to quantitative analysis of cooperativity for multiple binding sites of a single protein (although it can provide qualitative information on heteromultimeric binding) [30-32]. However, these practical approaches have not been integrated fully into the theoretical framework used for other substrate-ligand interactions. There are many challenges to elucidating cooperative interactions among transcription factors. Conditions in nuclei are difficult to measure and reproduce. All the protein domains that affect function must be included, sometimes making protein purification difficult. Nucleotide sequence flanking the core binding motifs that are sufficient to capture any effects that they may have on binding site shape should also be included [33-35]. Additional complexities are involved when relating *in vitro* binding data to transcriptional readouts. For example, cooperative interactions involving chromatin templates, which may occur through cooperative displacement and modification of nucleosomes, are typically not measured in such studies. Despite the inherent limitations with *in vitro* systems, computational and bioinformatic approaches have had some success in describing cooperativity (see e.g. [24, 36]) using high-throughput methods. Such approaches often continue to rely on previously analyzed, cooperating proteins to entrain the system and provide meaningful quantitative output [37].

Models oriented toward allosteric binding such as MWC can quantify the various conformers of the complex based on the free (unbound) concentrations of each ligand [4,5]. This is useful when measuring free ligand is relatively straightforward, as for the partial pressure of oxygen in the hemoglobin system or the concentration of Ca^2+^ in the calmodulin system. However, the analysis of transcriptional regulation may involve the measurement of one or more cooperative complexes as a function of total added ligand. For this context, where free ligand can be very difficult to measure, quantitative, experimentally accessible models have not been fully developed. Further, saturation binding may require high ligand concentrations, which are attainable for ligands like O_2_ and Ca^2+^, but can result in aggregation and precipitation of protein ligands. When the binding substrate is DNA or RNA, high ligand concentration may also produce significant off-target (i.e., non-specific) binding. Finally, saturation binding measurements can require the production of large quantities of purified protein, which can be a technical limitation.

Here, we revisit the use of Hill plots to quantify cooperativity, and illustrate key shortcomings as a motivation to develop a more practical and descriptive approach. We develop this approach within a broad theoretical framework applicable to any system involving multiple ligands binding to two or more distinct sites on a substrate, while focusing on the practical application of cooperative binding to DNA by proteins. In that context, we develop methods for quantifying cooperativity by two ligands binding to distinct sites, first measuring individual binding constants and then a cooperativity factor (previously referred to as the cooperativity parameter in, e.g., [22]). This provides the means to predict the binding behavior of cooperating proteins over the full range of concentrations. Motivated by recent identification and analysis of cooperative binding sites for Engrailed with its partner complex Extradenticle/Homothorax (Exd/Hth) [38], we devise a novel method for determining binding constants for individual ligands using competition assays [39], which we show has significant advantages over saturation binding assays. We go on to devise a method to find the cooperativity factor and the second equilibrium constant when only one of the two equilibrium constants can be accurately measured using single-ligand binding. Finally, because the cooperativity factor is thermodynamically meaningful, once it has been determined for pairwise interactions, we show how it can be employed to model more complex systems involving multiple components.

## 2. MATERIALS & METHODS

All computational methods used here broadly follow the same three steps, with different parameters and inputs. First, a binding system is defined with equilibrium constants and total concentrations of each component. Second, the equations relating these equilibrium constants and total concentrations to concentrations of individual species are specified. Third, these equations are manipulated and combined, either manually or using a computer algebra system (Wolfram Mathematica 10.4.0), to relate the desired quantities. Lastly, numerical values are substituted for parameters and the desired outputs computed and/or graphed. The graphs for Figs. 1-3 were produced using the Mathematica notebooks included as supplements. Microsoft Excel and Pacific Tech Graphing Calculator 4.0 were used to generate graphs for Fig. 4. Inkscape and Adobe Photoshop were used to compose figures.

In Fig. 1, the Hill plots are derived for the system shown in Fig. 1A. Using the equilibrium equations (Fig. 1A, lower) and conservation of mass equations, an expression for the total fractional occupancy as a function of free ligand concentration ([*a*]) and the parameters *K_A_*, *K_B_* and *n* was derived (manually in Fig. 1A of Peacock and Jaynes [2]; using Mathematica in Fig_1.nb within Supplemental_Mathematica_notebooks). By definition, a Hill plot shows *θ*/(1 – *θ*) as a function of [*a*] on a log-log plot, producing Fig. 1B. The maximum slope was calculated by taking the derivative at the point of 1/2 occupancy (always appropriate for 2 sites, see below).

In Fig. 2, a more complex system is described with two distinct ligands, *a* and *b*. Again using the relevant equilibrium and conservation equations, an equation describing the concentration of ternary complex, [*AaBb*], as a function of total ligand concentration [*a*]_T_ and parameters was derived (by hand in Fig. 2A of Peacock and Jaynes [2]; using Mathematica in Fig_2.nb within Supplemental_Mathematica_notebooks). Illustrative parameter values were selected to demonstrate the relevant concepts, and each curve calculated and plotted. Points of half-maximum occupancy were found by calculating the limit of [*AaBb*] as [*a*]_T_ approaches positive infinity (see Fig. 2B in Peacock and Jaynes [2]).

**Fig. 2.**
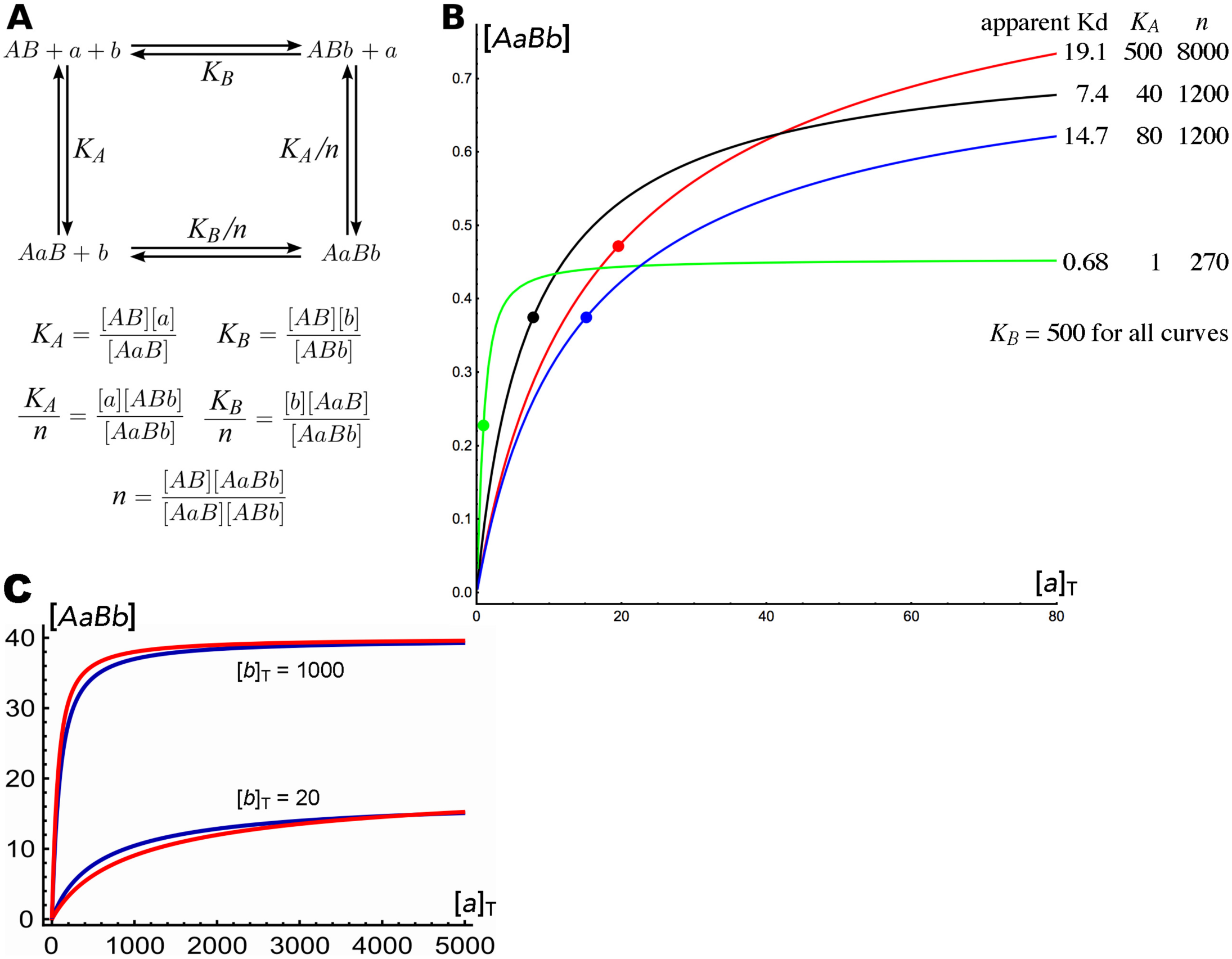
Two cooperating proteins binding to two different sites. <u>**A:** schematic of binding equilibria.</u> Either protein can bind first. The upper path from left to right represents initial binding by protein *b.* The equilibrium concentration of the *ABb* single-protein complex is governed by its Kd, *K_B_.* It can then bind protein *a* to form the ternary complex *AaBb.* The alternative pathway to the ternary complex is similarly diagrammed on the left. As the definitions of the various dissociation constants below show, not all 4 are independent. If we divide *K_B_* by the Kd that governs the dissociation of *b* from *AaBb*, we get the same quantity that we get if we divide *K_A_* by the Kd that governs the dissociation of *a* from *AaBb.* It is therefore convenient to define this ratio as the cooperativity factor *n*. <u>**B:** graphs of [*AaBb*] as a function of increasing [*a*]_T_, holding [*b*]_T_ constant.</u> For all graphs, [*AB*]_T_ = 1, [*b*]_T_ = 2, and *K_B_* = 500, while *K_A_*’S and cooperativity factors vary. The apparent Kd (based on a single-site model, see Fig. 2B in Peacock and Jaynes [2]) is the [*a*] at the point of half-maximal [*AaBb*], which is marked by a dot for each curve. Note that the relative amounts of ternary complex depend strongly on [*a*]_T_. The green curve, which has the lowest apparent Kd (0.68), actually shows the lowest [*AaBb*] at high [*a*]_T_. This is due to its relatively low *n*, which determines the [*AaBb*] at saturation with protein *a*, independent of *K_A_.* The black curve crosses the red curve, and also shows less binding at high [*a*]_T_ due to a lower *n*. The blue curve does not cross the red curve, and has a lower [*AaBb*] at all values of [*a*]_T_, despite having a lower apparent Kd! So, a ranking of apparent Kd’s from this type of experiment is not predictive of relative ternary complex formation overall. Derivations of expressions relating [*AaBb*] to [*a*]_T_ (and to [*a*], [*AaB*], and [*AB*]) are given in Fig. 2A of Peacock and Jaynes [2]. Derivations of expressions for [*AaBb*]max and for finding the apparent Kd are given in Fig. 2B of Peacock and Jaynes [2]. <u>**C:** relative ternary complex formation can be qualitatively different depending on the fixed [*b*]_T_ chosen for the experiment.</u> The two pairs of curves (upper and lower) represent [*AaBb*] formed on two sites (red and dark blue) as a function of increasing [*a*]_T_, differing only in the fixed [*b*]_T_. Note that at the lower [*b*]_T_, the site represented by the dark blue curve (*K_A_* = 5975, *K_B_* = 400, *n* = 113) forms more ternary complex throughout most of the experimental range, while at the higher [*b*]_T_, the site represented by the red curve (*K_A_* = 90,000, *K_B_* = 2808, *n* = 7000) forms more over the entire range. This illustrates another limitation of modeling cooperative binding using a single parameter. **NOTE:** Concentration units are not specified, because in all cases, these units (which includes the concentrations of ligands and substrate, as well as Kd’s) can be factored out of the governing equations, and do not affect the shapes of curves, or any of the conclusions.

In Fig. 3A,B, a competition system is treated, consisting of the Fig. 2 system plus unlabeled competitor, *U_AB_*, with its own cooperativity and equilibrium parameters. This system is used to describe 3 experiments [38], each with the same labeled DNA oligo (B1a), competing with an unlabeled oligo, either A2a, B1b, or B1a itself. Each experiment uses the same set of equations, but with different parameter values, chosen to approximate a situation encountered experimentally [38]. For Fig. 3A, the equations were solved to find [*AaBb*] as a function of added competitor [*U_AB_*]_T_. For Fig. 3B, the concentrations of complexes formed on the unlabeled competitor are solved for as a function of [*U_AB_*]_T_. In Fig. 3C, the three DNAs are considered without competitor, with fixed concentrations of both DNA and ligand *b*, and with increasing [*a*]_T_. This system is identical to that of Fig. 2, here solved for [*AaB*], [*ABb*] and [*AaBb*] in terms of [*a*]_T_. For complete methods, see Fig_3.nb within Supplemental_Mathematica_notebooks.

**Fig. 3.**
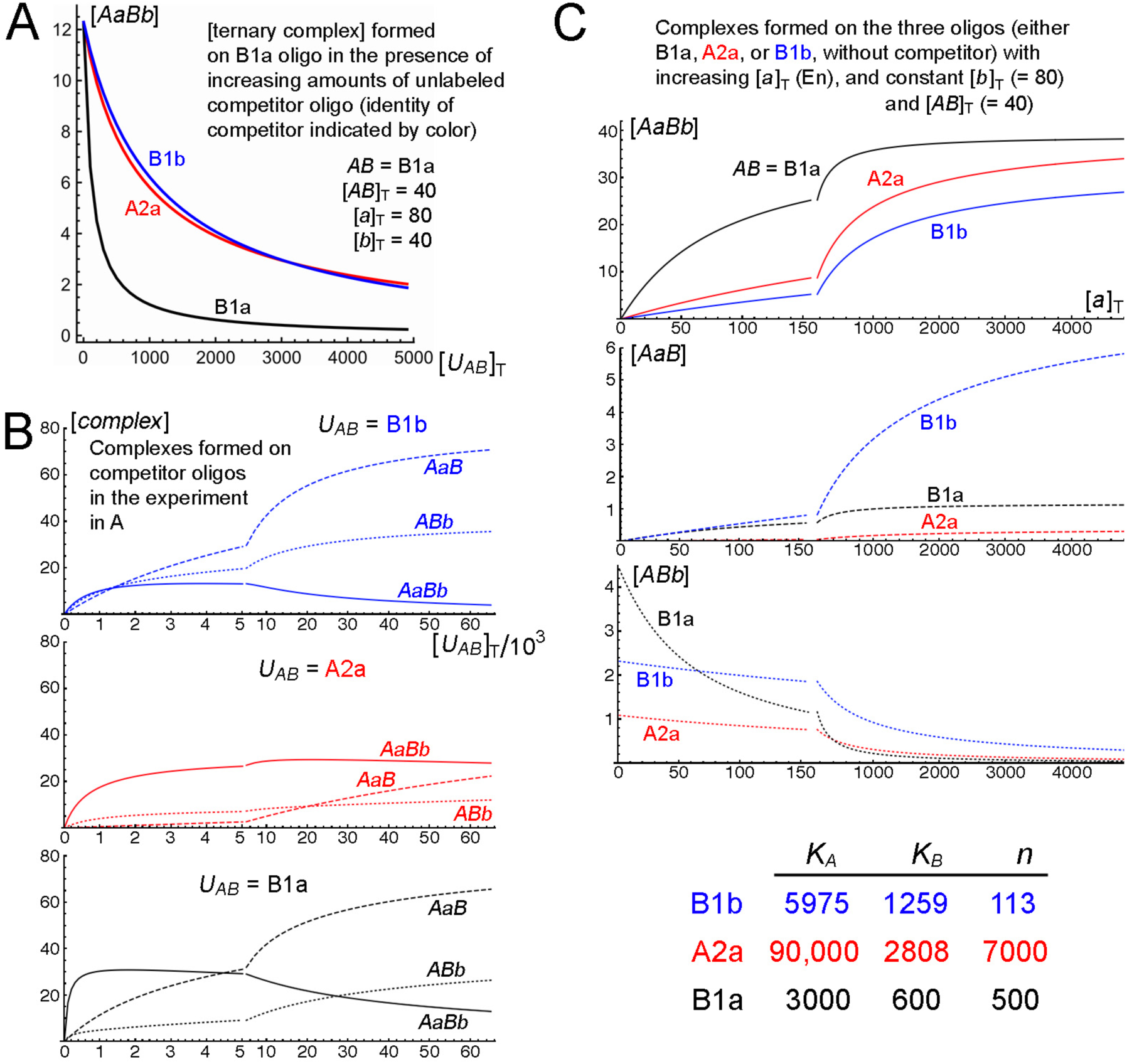
Competition curves measuring ternary complex have limited predictive power. Concentrations and Kd’s are in nM. <u>**A:** Ternary complex as a function of competitor. For each curve, the [*AaBb*], labeled ternary complex, is graphed as a function of an unlabeled competitor.</u> For each curve, the [*AaBb*], labeled ternary complex, is graphed as a function of an unlabeled competitor. The lower (black) curve is self-competition by the high-affinity site (B1a [38]), where the unlabeled competitor, *U_AB_*, is the same DNA sequence as the labeled binding site, *AB*. The other two curves show competition with two binding site oligos that have very different Kd’s and cooperativity factors (*n*), yet compete similarly for ternary complex formation by labeled B1a oligo. Note that self-competition is much more effective at all concentrations shown than is competition by either of the other oligos, while the other two oligos compete very similarly over a wide concentration range. <u>**B:** Forms of competitor oligo in the competition experiment.</u> Each graph shows the 3 bound forms for one of the competitor oligos. In each case, the solid line shows [*AaBb*], the dashed line shows [*AaB*], and the dotted line shows [*ABb*]. The upper panel shows the less cooperative low-affinity site (B1b, blue), the middle panel shows the more cooperative low-affinity site (A2a, red), and the lower panel shows the high-affinity site (B1a, black]. The left and right sections of each curve show two different ranges of [*a*]_T_, on two different scales. Note that at the higher concentrations of competitor, B1b forms mostly single-protein complexes, while A2a forms mostly ternary complex, reflecting its much higher cooperativity. At concentrations well beyond the range shown, all of each protein is incorporated into single-protein complexes, as the proteins are distributed over a vast excess of oligo. For oligo B1b (blue), we see the approach to this limit, while for oligo A2a (red), this approach is beyond the range shown. <u>**C:** Concentrations of complexes as a function of [*a*]_T_, holding [*b*]_T_ constant (no competitor).</u> Using the same oligos as in A (and B), the concentrations of the various protein-containing forms are graphed for a similar experiment as in Fig. 2. The left and right sections of each curve show two different ranges of [*a*]_T_, on two different scales. The top graph shows [*AaBb*] for each oligo, color-coded as in A and B. Note that despite the similarity of the blue and red competition curves in A, the oligo with the higher value of *n* (red) forms more ternary complex at all [*a*]_T_, and the red curve approaches the black curve at high [*a*]_T_. This provides a plausible explanation for the *in vivo* behavior of the binding sites represented by the blue and red curve: the one with the higher *n* (red) is more potent. It is more similar to the black curve than to the blue curve at high [*a*]_T_, suggesting that the ability to form ternary complexes at high [*a*]_T_ may explain the relative functionality of these binding sites *in vivo.* The middle and bottom graphs show [*AaB*] (dashed) and [*ABb*] (dotted), respectively, for each oligo, color coded as above. As seen in the competition experiment in B, the less cooperative oligo forms more binary complexes (blue) than does the more cooperative oligo (red), especially *AaB* at high [*a*]_T_, due to its having a lower Kd for binding each of the proteins. A similar phenomenon occurs at high concentrations of these oligos in the competition experiment: the less cooperative site sequesters more of each protein individually, while it forms less ternary complex than does the more cooperative site. These complexes are invisible in a competition assay, because the competitor oligo is unlabeled. For derivations of equations that can be used to generate these graphs, see Fig. 3A in Peacock and Jaynes [2]. For derivations of equations for graphing the total occupancy by each protein as a function of [*a*]_T_, see Fig. 3B in Peacock and Jaynes [2].

In Fig. 4A, a simple competition system is considered, consisting of a substrate DNA oligo (*A*) binding a ligand (*a*), in competition with an identical unlabeled DNA, *U_A_.* Solving the governing equations of this system, [*Aa*] is plotted as a function of [*U_A_*]_T_ for a given set of parameters (see also Fig. 4A in Peacock and Jaynes [2]).

**Fig. 4.**
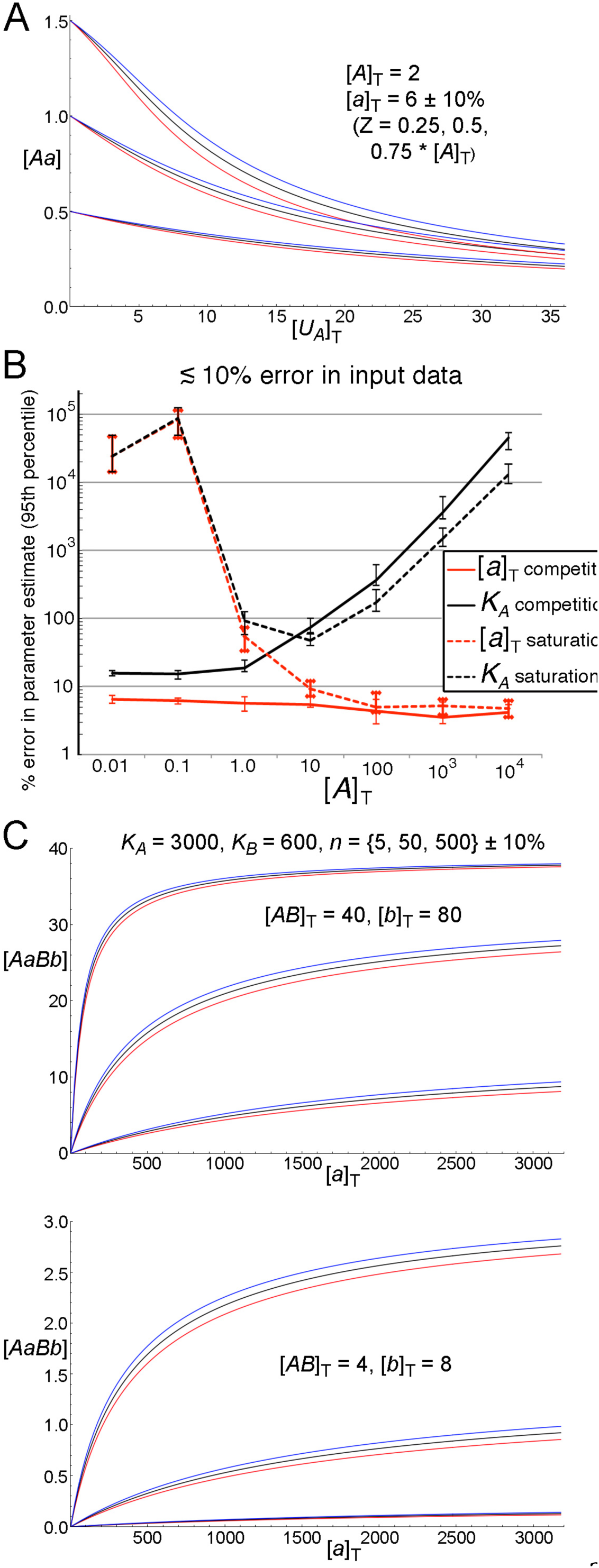
Illustrations of curve-fitting equations and error comparison. <u>**A:** Families of competition curves with different values of [*a*]_T_ and *K_A_.*</u> Labeled binary complex, [*Aa*], is graphed as a function of increasing total unlabeled binding site, [*U_A_*]_T_, with constant amounts of both labeled binding site, [*A*]_T_, and total ligand, [*a*]_T_. The applicable formula is [*U_A_*]_T_ = [*A*]_T_ ^∗^([*a*]_T_/[*Aa*] – *K_A_*/([*A*]_T_ – [*Aa*]) – 1). Sets of data points {([*U_A_*]_T_, [*Aa*])} along with known [*A*]_T_ can be used to find both *K_A_* and [*a*]_T_ as parameters using freely available curve fitting software (see text). The values used here are: [*A*]_T_ = 2 for all curves; [*a*]_T_ = 6, 5.4, and 6.6, and *K_A_* = {1.5, 5, 16.5}, {1.3, 4.4, 14.7}, {1.7, 5.6, 18.3}, for the black, red, and blue curves, respectively. The values of *K_A_* were adjusted to give 3 sets of 3 curves each with the same 3 initial values (without competitor), 0.5, 1.0, and 1.5. Note that each set of curves with the same starting value diverges significantly as competitor increases. <u>**B:** Performance of competition and saturation binding methods for simultaneously finding [*a*]_T_ and *K_A_.*</u> Monte Carlo analysis (100 trials are represented by each data point) of the accuracy of curve fitting to find both [*a*]_T_ and *K_A_* as parameters was run with 15- point data sets at 7 different [*A*]_T_ using either competition (fixed [*a*]_T_, varying [*U_A_*]_T_, as illustrated in A) or standard saturation binding (varying [*a*]_T_, no competitor). Data sets were generated by introducing random errors into calculated values of [*Aa*]. These errors were randomly drawn from a normal distribution (centered on zero) such that 95% of the errors were within ±10% of the actual value (standard deviation = 5%, mean error = 4.0%, median error = 3.4%). Percent errors are shown in the values found for each parameter ([*a*]_T_ and *K_A_*) using least-squares non-linear regression. These percent errors were ranked by increasing absolute value, and the 95^th^ largest (out of 100) plotted, with errors bars extending between the 90^th^ and 99^th^ largest. These error bars represent a 95% confidence interval for the true value of the 95^th^ error percentile, based on standard statistical analysis. Note that the best estimate for *K_A_* is provided by the competition method at low [A]_T_, which simultaneously provides a precise estimate for [_a_]_T_. See text for further explanation. <u>**C:** Family of curves with different values of *n*.</u> [*AaBb*] is graphed as a function of [*a*]_T_, holding constant [*b*]_T_ and [*AB*]_T_, for 3 different values of the cooperativity factor (*n*). *K_A_* = 3000, *K_B_* = 600, *n* = {5, 50, 500} for the black curves, *n* = {4.5, 45, 450} for the red curves, and *n* = {5.5, 55, 550} for the blue curves. The uppermost black curve corresponds to the black curve in Fig. 3C, top. Once *K_A_*, *K_B_*, and [*b*]_T_ are determined, the formula used to draw these curves can be used to find *n* from sets of data points {([*AaBb*], [*a*]_T_)} using freely available software (see text). The formula is

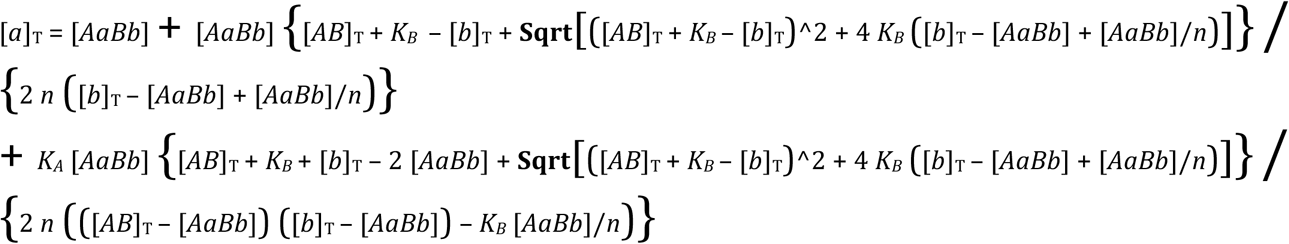 Note that the upper set of curves in the upper graph (which differ among themselves only by a change in *n* of 10%, like each set of 3 closely situated curves) are very close together, making it difficult to determine this *n* (= 500) using these values of [*AB*]_T_ and [*b*]_T_. However, when both are reduced by a factor of 10 (as shown in the lower graph), the upper set of curves (again representing *n* = 500) diverge more. Thus, *n* values that result in saturation of the probe (*AB*) can be more precisely determined by reducing its concentration, along with that of the fixed [*b*]_T_ (which is optimal for determining *n* when it is similar to [*AB*]).

In Fig. 4B, the results of competition and saturation binding experiments were computationally simulated under 7 sets of parameter values ([4]_T_, *K_A_*, and [*a*]_T_). To simulate a competition experiment, the system of Fig. 4A was used, and a series of 15 points ([*U_A_*]_T_, [*Aa*]) were calculated, with simulated experimental noise added to [*Aa*]. For the saturation binding experiment, the simple system of ligand *a* binding substrate *A* to form complex *Aa* was simulated to produce points ([*a*]_T_, [*Aa*]). For comparison with competition binding, it was assumed that [*a*]_T_ was actually a dilution series from an unknown stock [*a*]_T0_. The points generated in each simulated experiment were then fit with least squares regression [40, 41] to the corresponding equations used to generate them, but now with [*a*]_T0_ and *K_A_* unknown, and experimental noise in [*Aa*]. This simulation was repeated 100 times to produce a Monte Carlo estimate of the 95^th^ percentile of the absolute percent error of the parameter estimates, which is shown for each of the 7 conditions. For details, see Fig. 4B in Peacock and Jaynes [2], which shows the same system as Fig. 4B, again solved for ternary complex [*AaBb*] as a function of [*a*]_T_, plotted at a variety of parameter values, and showing the 50^th^ percentile as well as the 95^th^.

In Fig. 4C, the system of Fig. 2 is used to draw graphs for different values of the cooperativity factor *n*. This is done to demonstrate how the expression derived for [*AaBb*] as a function of total ligand [*a*]_T_ can be used to determine *n*, via regression analysis, once *K_A_* and *K_B_* have been determined (using, e.g., the method illustrated in Fig. 4A).

## 3. RESULTS AND DISCUSSION

### 3.1 One ligand binding cooperatively to two sites

In order to develop a practical, systematic approach to the problem of cooperative binding, we begin with the simplest case. A straightforward model to describe the binding of a single ligand to two sites is illustrated in Fig. 1A. A species with two distinct ligand binding sites (*AB*) can form a ternary complex (*AaBa*) with free ligand (*a*) by first forming either of two single-ligand intermediates, *AaB* or *ABa*. Adding a second ligand then converts either of these to the ternary complex. Assigning standard equilibrium binding constants (dissociation constants, or Kd’s) to each of these reactions gives the relationships shown in Fig. 1A below the flow chart. While there are 4 distinct Kd’s, one of them is not independent, but is completely determined if the other 3 are fixed. Reflecting this, we use only 3 variables to describe these Kd’s, noting that the two Kd’s governing the occupancy of site *A* are related to each other by the same factor that relates the two Kd’s that govern the occupancy of site *B.* We call this factor *n*, the cooperativity factor. It represents the fold decrease in Kd that results from prior occupancy of the other site, or, equivalently, the fold increase in affinity for a second ligand caused by binding of the first ligand. Thermodynamically, it is related to the change in free energy of association (or dissociation) of one ligand that results from binding by the other. A cooperativity factor greater than 1 indicates positive cooperativity, where bound ligand increases the affinity for additional ligands, while a cooperativity factor less than one means that bound ligand reduces the affinity for additional ligands. Using these relationships, for any specified Kd’s *K_A_* and *K_B_*, cooperativity factor *n*, and total concentration of *AB*, we can calculate the occupancy of each site, and the overall fractional occupancy, *θ*, at any concentration of ligand. The fractional occupancy as a function of free ligand concentration (both on a log scale) can then be displayed as a Hill plot [1, 42, 43], to see how such plots vary for different sets of constants.

Fig. 1B shows several interesting cases that illustrate some of the difficulties with using such plots for measuring cooperativity experimentally. The blue curves show two cases that conform to the situation of equivalent sites (*K_A_* = *K_B_*, or *K_A_*/*K_B_* = 1), for which the Hill formalism was originally derived, while the solid blue curve shows the classical case of positive cooperativity. The maximum slope (the purple line is tangent to the curve at the point of maximum slope) is greater than that without cooperativity, where the plot is a straight line of slope 1 (not shown). As is well known, this maximum slope approaches the number of cooperating sites (in this case 2) as the cooperativity becomes very large. When the cooperativity is negative (*n* < 1; *n* = 0.04 for the dashed blue curve), the plot shows the opposite shape, having a slope < 1 at its minimum point. (This maximum or minimum slope always occurs at 50% occupancy for two sites, discussed further below.) These effects depend strongly on the assumption of equivalent sites, a condition that is not often achieved with biological molecules, particularly for protein binding sites on DNA. When the two sites have different Kd’s and no cooperativity (*n* = 1), the Hill plot shows the same behavior as with equivalent sites and negative cooperativity, illustrated for the case where the Kd’s differ by a factor of 100 (Fig. 1B, dashed red curve).

Strikingly, positive cooperativity can be completely masked for non-equivalent sites [7], resulting in a straight line (solid red), which is indistinguishable from equivalent sites with no cooperativity (not shown). This, of course, occurs only for particular values of relative Kd and cooperativity, but in all cases of non-equivalent sites, apparent cooperativity is reduced. If the ratio of Kd’s is far enough from 1, it will look like negative cooperativity rather than positive cooperativity (not shown). The precise conditions for these effects are given in the legend of Fig. 1B, and derived in Fig. 1A of Peacock and Jaynes [2]. Clearly, then, in cases where binding sites may not be equivalent, Hill plots are highly unreliable indicators of cooperativity [44, 45].

### 3.2 One ligand binding cooperatively to two or more equivalent sites

The Hill equation assumes an implausibly high-order reaction mechanism, equivalent to the simultaneous binding of multiple ligands. However, in the special case of equivalent sites and very high cooperativity, the Hill formalism can serve as a good approximation, especially if the cooperativity occurs in a single step. Fig. 1C,D in Peacock and Jaynes [2] provides an illustration of Hill plots for this case, and for the related case of progressive cooperativity, where each additional bound ligand changes the affinity for subsequent ligands in equal increments.

To summarize the lessons that can be gleaned from these examples (see Fig. 1 in Peacock and Jaynes [2] for a full description), for a Hill plot to reveal either the number of cooperating sites or the degree of cooperativity, accurate binding data must be obtained for a wide concentration range in order to determine the point of maximum slope. This is because the maximum slope may not occur at 50% occupancy, and slopes at either extreme of concentration approach 1. Perhaps most unrealistically, these approaches are only effective for equivalent ligand binding sites, which is rarely expected for natural DNA binding sites and ligands.

### 3.3 Two proteins binding cooperatively to two sites: an alternative to measuring co-complex formation as a function of one protein concentration

If the Hill formalism is not appropriate for most realistic situations involving cooperative binding, what is the solution? The one clear advantage of the Hill equation is its simplicity, and this is of course also the source of its limitations. To better model the complexities of real life, it is useful to first study a relatively simple case in some detail, and then use the results to build up to more complex situations. We therefore consider the case of two ligands that cooperate on two distinct binding sites (here and in Sections 3.4 and 3.5). We will use the example of DNA binding proteins cooperating on nearby DNA sites as our model system, but the methods we describe are general. These methods apply to any case where two different ligands bind two distinct sites on a receptor or substrate, with cooperative interactions affecting the relevant binding constants.

This situation has been studied in some detail in a variety of contexts, and it is useful to consider commonly used methods and their limitations. One approach models the system as a single ligand binding to its site, by measuring an apparent Kd of one protein binding in the presence of the other (e.g. [46]). Conceptually, the single “site” is represented by the combination of binding sites and the ligand with constant concentration. Experimentally, we choose a fixed concentration of double-stranded DNA oligonucleotide (oligo) containing the cooperating pair of binding sites, along with a fixed concentration of one of the proteins, and measure the amounts of ternary complex that form as the concentration of the second protein is varied. Typically, the fixed concentrations are chosen based on preliminary experiments that reveal cooperative binding. When a chosen concentration of each protein alone gives little or no detectable binding to the oligo, while mixing all three components results in a clearly detectable ternary complex, positive cooperativity is indicated. It is often of interest in such cases to compare the affinities of related pairs of proteins for the same DNA sites, or of similar (e.g., mutated) sites for the same pair of proteins. We consider here and in Section 3.4 the methods typically used for such comparisons, along with their limitations. We use an example from the literature in some detail, to illustrate how a more complete analysis of the binding parameters can reveal important aspects of the underlying mechanism of binding.

First, we consider the option of measuring an apparent Kd as described above, and using it to compare two related sets of proteins or sites. A binding scheme to describe the situation is shown in Fig. 2A. It is very similar to that considered in Fig. 1A for a single ligand binding cooperatively to two different sites, but here there are two ligands, each binding to only one of the sites. As before, we assume that the binding can occur to each site independently (binding is not ordered), which implies that the ternary complex can dissociate in either of two ways. Also as before, the binding can be described fully using three parameters: *n*, along with the two binary complex Kd’s, *K_A_* and *K_B_.* In principle, *K_A_* and *K_B_* can be measured independently and directly, assuming that non-specific binding (including binding of each protein to the other site) occurs only with orders of magnitude lower affinity, which is often found to be the case. As we demonstrate later, once the individual Kd’s are known, *n* can be readily determined, at least in principle. In practice, there are interesting cases where one of the individual Kd’s is so high (i.e., the affinity is so low) as to be difficult to measure directly. In Section 5 of Peacock and Jaynes [2], it is shown how to leverage an individual measurement of only one of the Kd’s, along with the results of cooperative binding, to determine all three parameters.

Strikingly, in the situation described above where total concentration of protein *a* ([*a*]_T_) is varied, and total concentrations of both oligo ([*AB*]_T_) and protein *b* ([*b*]_T_) are fixed, a wide variety of binding behaviors can result. When comparing the concentration of ternary complex ([*AaBb*]) formed when using either of two sets of parameters, we find that [*AaBb*] for one set of parameters may be lower than for the other set at low [a]_T_, but higher at high [a]_T_. In other words, the curves of [*AaBb*] vs. [a]_T_ may cross (Fig. 2B). This can occur in either of two types of comparisons: comparing two different pairs of sites bound by the same two cooperating proteins, or comparing two sets of proteins binding to the same pair of binding sites. Importantly, in either case, if data from such an experiment are used to determine an apparent Kd for each binding curve, the expectation would be that where the apparent Kd is lower, [*AaBb*] would be higher at all values of [*a*]_T_. While this is, of course, true for a single ligand binding to a single site, this expectation fails with multiple ligands.

This is illustrated in Fig. 2B for the special case where *K_B_* is the same for all the curves. This would apply, for example, to comparisons of cooperative binding to an oligo by different members of a DNA-binding protein family, in combination with a common DNA-binding protein partner. The green curve gives the lowest apparent Kd, but it has the lowest [*AaBb*] at high values of [*a*]_T_. The red curve gives the highest apparent Kd, but it has the highest [*AaBb*] at high values of [*a*]_T_. And the blue curve gives a lower apparent Kd than does the red curve, yet it has a lower [*AaBb*] at all values of [*a*]_T_. The general reason for these “unexpected” outcomes is that the actual 3-parameter system gives more complex curves than can be modeled using a single parameter. The specific reason is that the concentration at which *AB* is saturated with ligand, which occurs at very high [*a*]_T_, can be very different for the different curves. For curves where *K_B_* is the same, as in Fig. 2B, the curve with the highest cooperativity factor has the highest saturation concentration of *AB*, regardless of *K_A_.* For example, the black and blue curves have different Kd’s but the same cooperativity factor, and therefore saturate at the same level ([*AaBb*] = 0.75; see Fig. 2B in Peacock and Jaynes [2] for the general formula). This is because at very high [*a*], *K_A_* becomes irrelevant. Given that both [*AB*]_T_ and [*b*]_T_ are constant, the amount of *b* incorporated into *AaBb*, in equilibrium with *b* and *AaB*, at very high [*a*] depends only on the Kd for dissociation of *b* from *AaBb*, namely *K_B_*/*n*.

From these examples, we can see that neither ranking the amounts of ternary complex formed at any one [*a*]_T_ nor measuring an apparent Kd in this type of experiment is predictive of relative complex formation overall. (The criteria used to define apparent Kd’s are given in Fig. 2B of Peacock and Jaynes [2].) Curves like these will cross each other under conditions that we can glean from the governing equations (given in Fig. 2A in Peacock and Jaynes [2] and described in Fig. 2C of Peacock and Jaynes [2]). Fig. 2C in Peacock and Jaynes [2] also gives an expression for the apparent relative affinity measured in SELEX-seq experiments of the type described in Riley et al., 2014 [47], in terms of the Kd’s and cooperativity factors of two cooperating ligands on two composite binding sites.

Another important issue when using this assay is illustrated in Fig. 2C. Two pairs of curves are shown, representing two different concentrations of the “fixed” ligand. As seen by comparing the two pairs of curves, one binding site forms more ternary complex at one fixed concentration, while the other does so at the other fixed concentration, throughout most of the concentration range of the varying ligand. That is, the relative binding behavior reverses in the two cases. The red curve represents a more cooperative site, and at higher concentrations beyond the range shown, it forms more ternary complex than does the less cooperative site, regardless of which concentration of fixed ligand is used.

Here, it is important to note that the formulae and concepts developed throughout this work depend only on relative, not absolute, concentration units. Therefore, we have not specified units for either concentrations or Kd’s, except for the specific example developed in Section 3.4 below. Furthermore, cooperativity factors are inherently unitless. Although the absolute nuclear concentrations of most transcription factors have not been established, they are generally thought to be in the range of nanomolar to micromolar [48]. This is also thought to be the concentration range for their functional set of binding sites, and for their individual Kd’s in binding to those sites. Therefore, it would be appropriate in applications involving these molecules to consider our values for concentrations and Kd’s to have units in the picomolar to nanomolar (nM) range.

Cooperativity for transcription factors has been quantified in very few cases. For λ phage *cro* and *c*I repressors, the measured interaction free energies correspond to cooperativity factors of <200 [49, 50]. (Each change in Kd of 10-fold corresponds to a change in interaction free energy of about 1.37 kcal/mol). Enhanceosomes in mammalian systems involve a number of cooperatively binding proteins [51-53]. Although in several of these cases, synergistic increases in site occupancy were attributed to cooperative binding, no quantitation of cooperativity was reported. However, interactions between transcription factors whose nuclear concentrations are below about 100 nM can result in cooperativity factors up to about 10^6^ without leading to much dimerization in solution. Thus, the cooperativity factors that we have used here are all physiologically plausible.

Clearly, in order to model the system accurately, we need to measure more than one parameter. In Section 3.5, we consider the commonly used method to measure individual Kd’s and compare it with a novel method, then go on to show how these can then be used to accurately determine the cooperativity factor *n*. Once these three parameters are known, we can predict the binding behavior, and specifically the relative amounts of each complex, at all concentrations of the components.

Before doing this, we consider another method commonly used to compare the affinities of different combinations of proteins and binding sites, competition experiments, where an unlabeled oligo competes with a labeled oligo for binding by fixed amounts of the proteins [54]. We will consider an example of this experiment in some detail, both to illustrate its limitations and to resolve an apparent paradox, leading to new biological insights.

### 3.4 A case study: insights from an analysis of competition assays

In a paper published in 2012 [38], Fujioka et al. characterized several cooperative binding sites for the Engrailed (En) protein within the *sloppy*-*paired* (slp) locus of Drosophila. En shows strongly cooperative binding to each of these sites with a cofactor complex, which contains one molecule each of the proteins Exd and Hth. This cofactor complex forms in solution and can be co-purified as a single complex when the proteins are co-expressed in bacteria. This stable complex has therefore been treated in binding studies as a single entity, Exd/Hth [35,37,46,54-57]. We compared the apparent affinities of four binding sites from *slp*, and found that two of them showed strong binding by En-Exd/Hth, while the other two were much weaker. These assessments were based on a combination of both direct binding assays and competition binding experiments like those described above. In all cases, ternary complex formation was monitored using gel shift analysis. Note that we refer to the complex containing En, Exd/Hth, and an oligo consisting of the cooperatively bound (composite) site as a ternary complex, because we model Exd/Hth as a single entity for the purposes of studying its cooperativity with En. Binding by the individual proteins was found to be relatively low, and in some cases was undetectable, consistent with a high degree of cooperativity in the binding. However, an apparent paradox was uncovered in that the two lower affinity sites showed distinctly different functional characteristics *in vivo*, despite very similar behaviors in the binding assays. The functional assay involved repression of a reporter transgene in En-expressing cells in the developing Drosophila embryo, which is dependent on a functional binding site for En and Exd/Hth [38]. We speculated that the subtle differences we observed in their apparent cooperativity might be responsible for their distinct functional potencies: the more cooperative site gave more complete repression *in vivo.* Despite this difference, which emerged from a limited set of direct binding assays, the competition assays, considered a good way to quantify relative affinities, showed no difference between them.

In modeling studies represented in Fig. 3, we revisit this issue by using the results from that work [38] to estimate individual Kd’s and cooperativity factors for these sites. We model the binding using these parameters, and make several noteworthy discoveries that may explain both the difference in function of the low-affinity sites and why the competition assays did not reveal a distinction between them. These results have important implications for the limitations of these assays, and provide guidelines for their effective use. We note that these results are based on published studies, and it is not our purpose here to establish new biological principles. Rather, we illustrate how quantifying both Kd’s and cooperativity factors allow exploration of binding behavior that is outside the concentration range used in the *in vitro* experiments themselves. This in turn can lead to new biological insights, including novel hypotheses that can be tested in subsequent studies. In particular, our analysis is not meant to either test or validate the assumption that Exd and Hth function strictly as a single unit when cooperating with En.

The competition binding assays yielded “competition curves” for each of the sites [38]. In this assay, as mentioned above, a labeled oligo containing the highest affinity site is bound by unlabeled proteins, with fixed concentrations. Increasing amounts of an unlabeled competitor oligo are added, which carries either the same sites as the labeled oligo or different sites, and the decrease in binding to the labeled oligo is quantified. Theoretical curves that closely match the published curves are presented in Fig. 3A for three of the sites. For simplicity, we use only one of the two high-affinity sites, which behaved similarly both in the binding studies and *in vivo.* To produce these curves, we used the limited set of direct binding studies available to estimate a range for the individual Kd’s and *n* for each site. We then refined these estimates to produce competition curves that match the data well. Although we do not consider these refined estimates to be precise, they nonetheless suggest a novel hypothesis as to why one of the sites functions better *in vivo.* More generally, the modeling results illustrate why the assays can be misleading, and provide guidance for their effective interpretation and use.

The main conclusions from these studies are 1) the two lower affinity sites have distinct binding behaviors that may explain their functional differences *in vivo*, 2) the competition binding studies did not reveal these differences because of the particular range of concentrations used, and 3) at those concentrations, the two lower affinity sites competed very similarly, but for different reasons. We now describe the results in some detail, to justify and fill out these conclusions. We then describe the lessons learned, one of which applies specifically to the interpretation of competition assays, while another reinforces the lesson from prior sections that in the face of the complexities of cooperative binding, even to two sites, it is necessary to measure multiple parameters in order to model the system effectively. This then provides the impetus to explore novel ways of measuring those parameters, presented in Section 3.5.

Fig. 3A shows that unlabeled oligos carrying the two low-affinity sites compete very similarly (red and blue) for binding to a labeled oligo carrying the high-affinity site. Of course, the high-affinity site itself competes much more effectively (black). Fig. 3B shows that the two low-affinity sites compete similarly for very different reasons, in terms of the complexes that they form as their concentrations increase. Although they each initially form more ternary complex (solid blue and red) than single-protein complexes, the oligo with the lower *n* (blue) binds relatively more of the single-protein complexes (dotted and dashed curves). This difference is magnified at higher concentrations, where the less cooperative oligo forms more of each of the single-protein complexes (blue dotted and dashed curves) than it does the ternary complex (solid blue), while the more cooperative oligo continues to form much more ternary complex (solid red). So, while the net result in the competition assay appears the same, this is specific to the choice of concentrations of labeled oligo and proteins. At other concentrations, differences would be more apparent. Rather than illustrating this, we show in Fig. 3C the differences in direct binding by these two oligos, which may explain their differences in function.

Fig. 3C, top, shows direct binding curves of the type in Fig. 2, [*AaBb*] vs. [*a*]_T_ with both [*b*]_T_ and [*AB*]_T_ constant. Here we see that more ternary complex is formed on the “red” site at all [*a*]_T_, compared to the blue curve. The more cooperative low-affinity site, which forms more ternary complex (solid red), is the one that functions better *in vivo.* In fact, its function *in vivo* is more like that of the high-affinity site than it is the other low-affinity site [38]. This may be explained by the fact that the amounts of ternary complex formed by the two higher-functioning sites (solid black and red) become more similar at high [*a*]_T_, and more distinctly different from the lower-functioning site (solid blue). That is, the red curve approaches the black curve, and separates from the blue curve, at high [*a*]_T_. This behavior is due to the higher cooperativity of the better-functioning site, as explained above for Fig. 2: the saturation value depends more on the cooperativity, while the relative behavior at low [*a*]_T_ is more dependent on *K_A_.*

Fig. 3C (middle and bottom graphs) illustrates the underlying reason for the “surprisingly strong showing” by the lower-functioning site in the competition assay. At a given set of concentrations, it forms more single-protein complexes (blue dashed and dotted) than does either of the other two sites (red and black). The results of a competition experiment depend on the total amount of each ligand bound by a site, rather than only the amount of ternary complex. Thus, less ternary complex is made up for by the formation of more single-protein complexes. This is again consistent with the lower cooperativity of the lower-functioning site (it forms relatively less ternary complex and more of the single-protein complexes).

In the competition assays, the total amount of unlabeled, competing binding site goes well beyond the range used in direct binding assays for the same site, typically up to hundreds of fold more. At such high concentrations, single protein complexes can dominate, even though very little of them form in direct binding assays. At the concentrations used in these experiments, this was the case. Therefore, only if sites are independently known to have either similar Kd’s for formation of each single-ligand complex, or to have similar cooperativity factors, can we expect competition assays to reveal a simple set of “relative affinities”.

As these examples emphasize, modeling the binding of cooperating ligands using a single parameter can only have predictive power for occupancies of sites over a limited concentration range. This limitation should be taken into account when interpreting experiments based on high-throughput methodologies. An example is given in Fig. 2C of Peacock and Jaynes [2].

### 3.5 A comprehensive method for modeling relative complex formation over a wide range of concentrations

The foregoing argues for measuring individual Kd’s and a cooperativity factor in order to model binding by a cooperating pair of ligands. To this end, we first describe a less well-known method that has significant advantages over standard methods for determining an individual Kd. We then show how the cooperativity factor can be determined once both individual Kd’s are known. In Section 5 of Peacock and Jaynes [2], this methodology is extended to find both the cooperativity factor and the second Kd when only one of the individual Kd’s can be accurately determined using single-ligand binding.

Standard methods have been described for determining individual Kd’s that involve simply measuring complex formation as a function of ligand concentration (the “saturation binding" method; see e.g. [41]). However, in cases where cooperativity is high, individual Kd’s can be challenging to determine accurately, particularly in cases where either the available amount of ligand or its tendency to aggregate precludes obtaining data at high ligand concentrations. Our alternative method uses competition assays, similar to those illustrated in Fig. 3A, except that only a single ligand is used to determine its individual Kd.

Using competition assays to determine an individual Kd has significant advantages, particularly in the case of DNA binding proteins. Preparation of proteins for binding assays often involves concentrating them in a way that can cause denaturation, and for this and other reasons, the fraction of protein that is active in binding may be difficult to determine. In such cases, directly measuring the amount of protein provides only an upper limit on its effective concentration. Using a competition binding assay, we can straightforwardly determine both the Kd and the active concentration of ligand from the same data set. We note in this context that recently developed high-throughput methods for comparing DNA affinities [58] do not always distinguish between absolute and active concentrations of ligand. Further, it is important that these high-throughput methods include exemplars for entrainment and validation that have been characterized by methods grounded in solution biochemistry, such as the following.

Fig. 4A shows examples of binding curves from modeling this type of competition experiment, in which constant total amounts of labeled oligo ([*A*]_T_) and ligand ([*a*]_T_) are used in combination with increasing amounts of unlabeled oligo ([*U_A_*]_T_), and the resulting ligand-substrate complex ([*Aa*]) is quantified. [*Aa*] decreases as [*U_A_*]_T_ is increased, while all other quantities are constant. [*A*]_T_ is known, and [*a*]_T_ and *K_A_* are determined as parameters in a non-linear regression analysis. Values for constants were chosen to illustrate why the approach can yield these two parameters independently. When any family of curves representing different Kd’s and ligand concentrations start at the same point (i.e., they give the same amount of complex without competitor oligo), they have significantly different shapes, and so diverge from their common starting point, no matter where that starting point is. The differences between the 3 colors is a 10% change in [*a*]_T_. The values of *K_A_* are adjusted for each curve to give the same initial value of [*Aa*]. This difference in shape can be captured during regression analysis to give the two parameters independently. Fig. 4A of Peacock and Jaynes [2] gives a derivation of the expressions used. These expressions can also be used in a high-throughput analysis to determine the relative affinities of related binding sites, where the highest affinity site is labeled, and measurements are made using a panel of unlabeled sites of how well they compete for binding to a fixed amount of protein, as described in Hallikas, et al., 2006 [59]. Fig. 4A of Peacock and Jaynes [2] gives an exact expression for determining the relative affinity in such an experiment, and also a simple approximation for the case where only a small fraction of labeled oligo is bound without competitor (which is different from that given in Hallikas et al. and has the correct limit behavior).

We tested the ability of this method to give precise values for the parameters *K_A_* and [*a*]_T_ under different conditions, and compared it to the traditional saturation binding method. First, a few words about the latter method. It is possible, in principle, to use saturation binding to determine both *K_A_* and [*a*]_T_ simultaneously. This can be done by varying [*a*]_T_ in a systematic way (for example by diluting a stock solution) and measuring the resulting [*Aa*], without initially knowing the actual values of [*a*]_T_. If we let the reference value of [*a*]_T_ be [*a*]_T0_, and each experimental value be [*a*]_T0_ / Δ (Δ is the “dilution factor”, which varies for each data point, while [*a*]_T0_ is a constant to be determined from regression analysis) then the relevant formula is Δ = [*a*]_T0_ / {[*Aa*] ^∗^ ({*K_A_* / ([*A*]_T_ – [*Aa*])} + l)}. From the data set {(Δ, [*Aa*])}, and knowing [*A*]_T_, we can use regression analysis to simultaneously find the two parameters *K_A_* and [*a*]_T0_. Although this can in principle work effectively, it requires using a set of experimental conditions that are unknowable before the Kd is determined. The optimal conditions are when [*A*]_T_ ~ 10^∗^*K_A_* (of course, *K_A_* is initially unknown). Even under these optimal conditions, data that are precise to within about 10% are required to determine *K_A_* within about 50% and [*a*]_T_ within about 10% (Fig. 4B).

In contrast, using the competition method described above, it is possible to use data accurate to within ~10% to determine *K_A_* within about 20% and [*a*]_T_ within about 7% (Fig. 4B). Importantly, the competition method provides this level of precision as long as the [*A*]_T_ used is less than or ~ *K_A_*. This contrasts with the saturation binding method, which becomes much less effective at determining *K_A_* when [*A*]_T_ is either above or below 10^∗^*K_A_* by several-fold or more. The flexibility and resolving power of the competition method allows us to define an optimal approach to precisely determining both *K_A_* and [*a*]_T_: use the lowest [*A*]_T_ that allows precise quantitation of [*Aa*], and the highest [*a*]_T_ available, up to a [*a*]_T_ that gives [*Aa*] ~ [*A*]_T_ / 2 without competitor *U_A_*. Data taken under these conditions, and with increasing *U_A_* so that [*Aa*] is reduced to 1/3 or less of its initial value without *U_A_*, will give the most precise value practicable for *K_A_*, along with a somewhat more precise value for [*a*]_T_.

These conclusions are put into context in Fig. 4B, which shows the results of a Monte Carlo analysis of the two methods. A random error was introduced into data sets, and regression analysis was used to find *K_A_* and [*a*]_T_ as parameters from the data. The resulting errors in the parameters are shown, as a function of the chosen value for [*A*]_T_. The value of *K_A_* was fixed at 1. This is justified by the fact that it is only the relative values of [*A*]_T_, [*a*]_T_, and *K_A_*, and not their absolute values, which determine the shapes of the curves, and therefore how precisely the parameters can be determined. As Fig. 4B illustrates, when [*A*]_T_ exceeds *K_A_* by more than 10-fold ([*A*]_T_ >10 in this case), determination of *K_A_* becomes increasingly unreliable (approaching or exceeding 100% error at least 5% of the time) with both methods. With [*A*]_T_ between *K_A_* and 10^∗^*K_A_*, both methods provide similarly reliable estimates of both parameters. Importantly, with [*A*]_T_ < *K_A_*, the competition method becomes more reliable, while the saturation method fails completely. These results support the strategy given in the previous paragraph for finding the two parameters simultaneously to the best possible precision under a wide variety of circumstances. Similar results are obtained with different errors in the input data sets (1% and 5%, Fig. 4B of Peacock and Jaynes [2]). Errors at the 50^th^ percentile in the error distribution (Fig. 4B of Peacock and Jaynes [2]) follow the same qualitative pattern as those at the 95^th^ percentile (Fig. 4B), illustrating that the overall error distribution is similar in all cases as a function of the chosen [*A*]_T_.

For both the competition and the saturation binding methods, a more precise value for [*a*]_T_ and a less precise value for *K_A_* are obtained when higher values of [*A*]_T_ are used above ~10^∗^*K_A_*. The precision for *K_A_* achievable within an experiment never exceeds that for [*a*]_T_. This is because in both formulae, *K_A_* is divided by ([*A*]_T_ – [*Aa*]), whereas [*a*]_T_ (or [*a*]_T0_) is divided by [*Aa*], which is typically lower when averaged over the data points than is ([*A*]_T_ – [*Aa*]). Therefore, a smaller change in [*a*]_T_ compensates for a larger change in *K_A_.* This causes the bounds placed on [*a*]_T_ by the data to be more stringent than those placed on *K_A_.*

For curve fitting to determine an individual protein’s Kd and its concentration by the above methods, we can use freely available software (e.g., at “statpages.org/nonlin.html”), along with the expressions given above and in Fig. 4A of Peacock and Jaynes [2] (which also includes their derivations). Fig. 4A of Peacock and Jaynes [2] also includes equations to provide initial guesses for the parameters from one or two data points, which are sometimes needed for the regression algorithm to converge.

It is often useful, from an experimental point of view, to include non-specific DNA in such binding experiments, to minimize the effects of contaminating DNA binding proteins that may come through the purification process. Therefore, we developed an alternative methodology which allows accurate determination of [*a*]_T_ and *K_A_*, as well as the Kd for non-specific binding, as parameters from curve fitting. The expressions used for this purpose, along with derivations and a description of how to analyze the data, are given in Fig. 4C of Peacock and Jaynes [2]. As an illustration of why this can work, if non-specific DNA is included in the experiment shown in Fig. 4A, the apparent Kd’s change without a significant change in the shapes of a curves, as long as the ratio of the non-specific Kd to each of the specific Kd’s is much greater than 1. So, the set of curves shown there is indistinguishable from that resulting from the inclusion of non-specific DNA with a Kd of 1000 at a concentration of 4000, with the same values for [*A*]_T_ (= 2), and [*a*]_T_ (= 6, 5.4, and 6.6), but with all of the *K_A_*’s reduced by a factor of 5.

Once individual Kd’s have been determined for both cooperating proteins, *n* for the composite site can be found readily. A simple method is illustrated in Fig. 4C. Here, the same equation used to generate curves in Figs. 3B,C and 4C is used to show how a 10% change in *n* affects the amount of ternary complex that forms as one ligand increases in concentration, while the total amount of the other is held constant. The same expression can be used in regression analysis to determine *n* from data of this kind. The upper panel of Fig. 4C shows 3 families of 3 curves each. Each family has a different value for n, and within each subfamily (those lying close together), this difference is 10%, either above (blue) or below (red) the middle value (black). The uppermost black curve in this graph uses *n* = 500, and it is the same as the black curve in Fig. 3C, upper panel. The other two subfamilies have *n* reduced by 10 and 100-fold, but use the same Kd’s. The relative amount of separation within each subfamily suggests how precisely *n* can be determined from data in such an experiment. The upper panel shows that the higher *n* value will be relatively difficult to determine accurately with the chosen concentrations of oligo and fixed protein. However, if we reduce these values, the curves for *n* = 500 ± 10% are more widely separated, as illustrated in the lower graph of Fig. 4C (both [*AB*]_T_ and [*b*]_T_ are reduced by 10-fold relative to the upper graph). In this case, it becomes easier to accurately determine *n* = 500, while it may become more difficult to determine the lower *n* values used here.

In order to get the most accurate determination of *n*, [*AB*]_T_ and [*b*]_T_ should be chosen to allow ternary complex formation to approach 50% saturation, but not exceed it. However, if this is not achievable with a [*AB*]_T_ that is high enough to allow accurate quantitation of ternary complex, it may be best to use the lowest [*AB*]_T_ (and [*b*]_T_, which should be comparable) that does allow accurate quantitation, and also results in less than 50% saturation at the highest achievable [*a*]_T_. The latter, of course, may be limited by the amount of available ligand. For regression analysis to determine *n*, we use the expression given in Fig. 4C (and derived in Fig. 2A of Peacock and Jaynes [2]).

The method described above requires that each individual Kd be measurable. However, for some ternary complexes, one of the ligands is not observed to bind alone, even at concentrations much higher than those required for strong, cooperative binding. A well-known example of this is described by Jin et al., 1999 [60], involving cooperative binding by the yeast transcription factors a1 and α2. Binding by α2 alone was seen, and addition of a1 resulted in much greater complex formation, suggestive of highly cooperative binding. However, even at the highest concentrations tested, no binding by a1 alone was observed. In order to extend our method to this type of situation, we devised equations for curve fitting to find both the Kd of the weakly binding ligand and the cooperativity factor from gel shift analyses similar to those presented by Jin et al. [60]. This method is described in Section 5 of Peacock and Jaynes [2].

### 3.6 Generalizing the methods to more than two binding sites and cooperating ligands

How can this methodology help us in understanding the functions of more complex composite binding sites, such as those often found in higher eukaryotic genes? Once the Kd’s and pairwise cooperativity factors have been determined for pairs of individual sites that make up the composite site, it is straightforward to model them as an interacting network of pairwise interactions. In a typical “Boltzmann” statistical thermodynamic version of such a model ([24] and references therein), the free energy differences induced by each pairwise interaction are combined to give a relative free energy, and therefore a relative occupancy under any specified conditions, for each possible complex. The Kd’s and cooperativity factor as defined above naturally feed into this type of model, because they can be readily associated with free energy differences among the various complexes. Although the behavior of such sites has been modeled without information about the pairwise interaction parameters (e.g., [24]), including those parameters would likely yield more meaningful, mechanistic models with wider predictive power that extends well beyond the range of the experimental data [61].

For DNA binding proteins in particular, it may be common for pairwise interactions to dominate the system, mediated by separable protein-DNA and protein-protein interaction domains. For example, in the cooperative interactions between λ phage *cro* and cI repressors, pair-wise interactions between nearest neighbors appear to dominate [49, 50]. In such cases, we can measure the individual protein-DNA Kd’s and pairwise cooperativity factors, to fully describe the behavior of complexes involving several DNA binding proteins. In these cases, the free energy of dissociation of each of the ligands from the 3-ligand complex can be accounted for by the dissociation energy of the two 2-ligand complexes which contain that ligand, so the system as a whole is solved by knowing each of the pairwise Kd’s and cooperativity factors. We now summarize the relationships among these quantities (which are measurable using the procedures described above) and the dissociation constants and cooperativity factor of a 3-ligand complex. Details are provided in Fig. 6 of Peacock and Jaynes [2].

We now need subscripts for each pairwise cooperativity factor to distinguish which pair of ligands it is associated with, as well as another cooperativity factor associated with the 3-ligand complex. This factor is an independent quantity in the general case where additional free energy (positive or negative) may be associated with the formation of the 3-ligand complex beyond that associated with the formation of each 2-ligand complex.

As derived for a 2-ligand complex containing *a* and *b* in Fig. 2A:

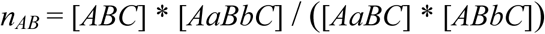

and, by analogy,

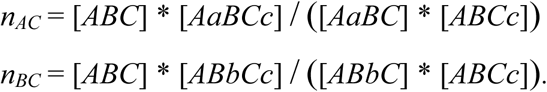

We can find *n_ABC_* to be:

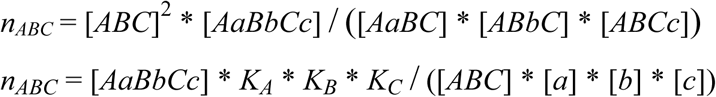

where

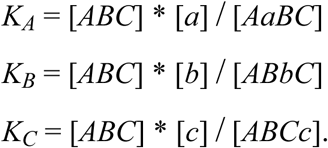

Now, how is the “new” cooperativity factor for the 3-ligand complex related to those of the 2-ligand complexes? Complete dissociation of the 3-ligand complex involves the sum of 3 free energies. For one possible dissociation route, these are: the free energy change when ligand *a* dissociates, that when ligand *b* dissociates from the 2-ligand complex, and that when ligand *c* dissociates from the single-ligand complex.

Because of the basic relationship between the standard Gibbs free energy and the Kd:

ΔG^0^ = – R ^∗^ T ^∗^ ln(Kd), where R is the gas constant and T is the absolute temperature, adding these 3 free energies is equivalent to multiplying together 3 dissociation constants: that governing the dissociation of ligand *a* from the 3-ligand complex, that governing the dissociation of ligand *b* from the 2-ligand complex containing ligands *b* and *c*, and that governing the dissociation of ligand *c* to release the free DNA.

This product is:

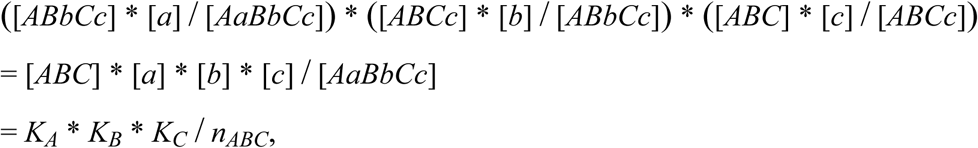

where the last equality comes from the last expression above for *n_ABC_*. So, 1 / *n_ABC_* represents the “extra” free energy in the complex that is due to cooperative binding; i.e., this “extra” ΔG^0^ = R ^∗^ T ^∗^ ln(*n_ABC_*). If there is no cooperativity, *n_ABC_* = 1, and the total free energy of dissociation is just the sum of those for each ligand individually. From the definitions above, we have, for the Kd that governs the dissociation of ligand *a* from the 3-ligand complex:

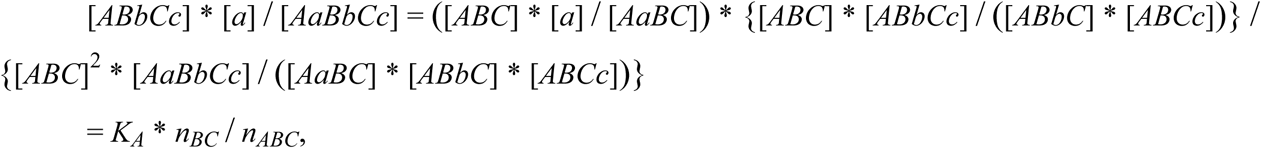

and similarly for the other dissociation constants from the 3-ligand complex.

Now, what do we expect for *n_ABC_* in the case where the free energies of interaction within the complex consist solely of those found within the respective 2-ligand complexes? In this case, as derived in Fig. 6 of Peacock and Jaynes [2],

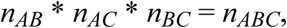

and we already have enough information from analysis of the 2-ligand complexes to predict the behavior of the entire 3-ligand system (see Fig. 6 of Peacock and Jaynes [2], which also provides a straightforward method for testing whether this relationship holds in any particular case).

We can follow an analogous procedure to characterize a 4-ligand system. If all of the interactions leading to cooperativity are contained within pairwise interaction domains that are not significantly affected by higher-order complex formation, then the sum of the free energies from the pairwise interactions equals the total cooperative free energy of the entire complex, and:

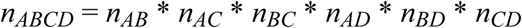

Generally, for *j* ligands binding to distinct sites on a substrate and cooperating solely through pairwise interactions, the cooperativity factor is the product of the

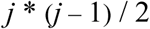

possible pairwise cooperativity factors, which can each be measured by studying the ternary complex containing those two ligands, using the methods given here, and in Peacock and Jaynes [2].

## 4. SUMMARY AND CONCLUSIONS

Most currently used methods for quantifying cooperative binding by transcription factors to DNA do not provide the means to accurately predict binding behavior over a wide range of concentrations. The Hill equation and Hill plots are only useful for quantifying cooperativity when binding sites are equivalent, which is rarely the case for DNA binding proteins, as they typically bind to a variety of sites with different affinities. Even in the rare cases where cooperating sites are equivalent, different modes of cooperativity are possible, and these have sufficiently different behaviors in Hill plots as to make quantifying the cooperativity difficult.

In recent years, two main approaches have been used to compare cooperative binding of either 1) two different proteins to the same sites or 2) the same two proteins to different sites. Both of these, while they provide some useful information, have serious limitations for predicting binding behavior over a wide concentration range. One involves holding the concentrations of both a binding site oligo and one protein constant while varying that of the second protein. We have shown that this can result in binding curves that cross, precluding the general usefulness of this approach in isolation to accurately model binding behaviors of cooperating proteins. However, this approach is useful for quantifying cooperativity once individual proteinbinding site Kd’s have been determined.

A second approach is to compare the ability of unlabeled oligos containing different sites to compete for binding (by two cooperating proteins) to a labeled binding site-containing oligo. We have shown that this approach, too, provides only partial information about the system and can therefore give misleading results. For example, two oligos can compete very similarly under one set of conditions, while the occupancies of these sites can be very different under different conditions.

This analysis of existing methods argues for a more comprehensive approach that can use experimental data obtained over a limited range of concentrations to predict binding behavior over the full concentration range. We therefore developed two new tools to achieve this end. The first tool is a new approach to determining individual protein-binding site Kd’s. Active protein concentrations must be determined in order to obtain accurate Kd’s, and our approach allows the simultaneous determination of both of these, as parameters in non-linear regression analysis, using data from oligo competition assays. We described an optimized approach to give maximum accuracy, which mandates using the lowest concentration of labeled oligo which allows robust quantitation, along with an amount of protein that gives around 50% occupancy. Holding both of these constant, increasing amounts of unlabeled oligo identical in sequence to the labeled oligo are added. Quantifying the resulting reduced binding to the labeled oligo gives a data set that is fed into freely available software (e.g., at “statpages.org/nonlin.html”), along with the equation we have derived. This can have very significant advantages over previously described methods that involve saturation binding (increasing concentrations of protein). One advantage is that our method typically requires lower protein concentrations, avoiding problems of precipitation or aggregation at high concentrations. The second advantage is that both active protein concentration and Kd can be accurately determined without prior knowledge of either parameter.

The second new tool is the means to extract, via regression analysis, the cooperativity factor for binding by a pair of proteins once the individual protein–site Kd’s have been determined. This involves holding the concentrations of a labeled oligo carrying the two cooperating binding sites and one of the proteins constant, while measuring cooperative complex formation as the other protein concentration is varied. Armed with the cooperativity factor and the two individual Kd’s, binding behavior can be predicted and compared over the full concentration range of each interacting species.

We also provide modifications to these methods that extend their applicability in special cases. First, in cases where contaminating DNA binding proteins co-purify with the specific protein being studied, it is often advantageous to include non-specific DNA in the experiments used to measure the Kd, which reduces significantly the inaccuracy that can result from these contaminating proteins competing for binding to the labeled oligo. We describe the method and give equations for regression analysis to find the specific Kd and the non-specific Kd along with the active protein concentration. Second, we provide a means to extract both Kd’s and the cooperativity factor (as parameters in regression analysis) for cases where only one Kd can be measured directly, which may occur when one cooperating protein binds very weakly on its own. This requires simultaneously measuring both ternary complex formation and the accompanying single-protein complex as the weakly binding protein’s concentration is varied.

We provide a summary of the main formulae used in this paper, along with their applications in the methodology developed here, in Fig. S1.

## 5. ACKNOWLEDGEMENTS

This research was supported by the National Institutes of Health, Awards R01GM050231 and R01GM117458 to JBJ. We give a special thanks to Miki Fujioka for the graphical abstract.

